# Sex dimorphism in the aged metabolic phenotype of smoothelin-like 1 (SMTNL1) deficient mice

**DOI:** 10.1101/2023.09.19.558520

**Authors:** Megha Murali, Nikolay Alabi, Prasanth K. Chelikani, Justin A. MacDonald

**Affiliations:** Department of Biochemistry & Molecular Biology, Cumming School of Medicine, University of Calgary, 3280 Hospital Drive NW, Calgary, AB, T3N 4Z6, Canada; Faculty of Veterinary Medicine, University of Calgary, 3330 Hospital Drive NW, Calgary, Alberta, T2N 4N1, Canada

## Abstract

Smoothelin-like 1 (SMTNL1) is expressed in smooth and skeletal muscle tissues as well as a variety of steroid hormone-sensitive tissues. SMTNL1 can play a sex-dependent regulatory role in skeletal muscle metabolism in mice. Previous studies have documented appreciable changes in muscle morphology and metabolic function of young male mice with genetic deletion of *Smtnl1*. SMTNL1 can also impact the energy metabolism and insulin sensitivity of female mice during pregnancy. Therefore, we investigated the metabolic outcome of global SMTNL1 knockout (KO) in male and female mice with advancing age using a comprehensive lab animal monitoring system (CLAMS). With ageing, body weight gain was markedly higher with a concomitant increase in whole body adiposity as well as specific white adipose depots in the absence of SMTNL1. Moreover, this genotypic difference in whole body adiposity was greater in the female cohort. The deletion of SMTNL1 was also associated with delayed satiety in mice fed a high fat diet, which was more pronounced in the female mice. A significant genotypic difference was also revealed for the metabolic energy balance in 12 month old animals of both sexes. The KO animals were metabolically less efficient and displayed a preference for carbohydrate catabolism. However, reduced glucose tolerance was observed only in the female group with the deletion of SMTNL1. Taken together, the current findings establish a novel role for SMTNL1 in modulating adiposity and energy metabolism with ageing in a sex dimorphic way.

## INTRODUCTION

Aging triggers complex metabolic and physiological changes that affect all organs and impacts on homeostatic communication among tissues (Guo et al. 2022; Finkel 2015; Palmer and Jensen 2022). Ageing is generally associated with a reduction in basal metabolism, energy expenditure during physical activity and in food intake (Finkel 2015). However, an increased accumulation of adiposity in middle-age can elevate the risk of metabolic dysregulation in old-age, with a change in fat deposition from subcutaneous to visceral. Other major metabolic impairments include uncontrolled hepatic gluconeogenesis, hyperinsulinemia, adipose lipogenesis, and defective glycogen and glucose handling in skeletal muscle (Barzilai et al. 2012). Many hallmarks of human aging (i.e., lymphocyte infiltration and tissue inflammation, stem cell exhaustion, decline in mitochondrial function, impaired proteostasis, insulin sensitivity and dysregulated nutrient sensing, increased adiposity, loss of muscle strength and motor function) are captured in murine models (Mitchell et al. 2016; Folgueras et al. 2018; Houtkooper et al. 2011; Huffman and Barzilai 2009). Proteomic surveys have characterized biomolecular changes that occur in different organs during aging (Petr et al. 2021) and revealed patterns of metabolic remodeling that characterize age-associated changes in mice coping with the increasing energetic cost to maintain health, energy, and redox balance during aging (Petr et al. 2021). *In vivo* studies in rodent models have also revealed that single genetic manipulations can ameliorate some age-related metabolic phenotypes.

Smoothelin-like 1 protein (SMTNL1) is found in smooth and skeletal muscle tissues as well as a variety of steroid hormone-sensitive cellular locales (Bodoor et al. 2011; Borman et al. 2004; Murali and MacDonald 2018; Wooldridge et al. 2008). SMTNL1 is regulated by post-translational modification (Borman et al. 2004; Wooldridge et al. 2008); namely, its phosphorylation by cyclic nucleotide dependent protein kinases can elicit changes in cellular locational targeting. Two known biological functions of SMTNL1 are reported: first, the protein can influence contractile properties of both smooth and skeletal muscle (Turner et al. 2019; Wooldridge et al. 2008) with interactions defined for the myosin-targeting subunit of the myosin phosphatase holoenzyme (Lontay et al. 2010; Borman et al. 2009), the actin thin-filament protein tropomyosin (Ulke-Lemee et al. 2010; MacDonald et al. 2012), and/or the intracellular calcium- receptor calmodulin (Ishida et al. 2008; Ulke-Lemee, Turner, and MacDonald 2015). Second, SMTNL1 can function as a transcriptional regulator of the progesterone receptor that ultimately provides phenotypic reprogramming of smooth and skeletal muscle beds (Bodoor et al. 2011; Lontay et al. 2015).

Regrettably, most preclinical metabolic research is conducted with male mice; however, a few investigations have defined important sexual dimorphism with respect to metabolic responses of aging male and female mice (Chaix et al. 2021). Indeed, SMTNL1 is known to play a sex-dependent regulatory role in skeletal muscle metabolism in mice. Previous studies have documented appreciable changes in muscle morphology and metabolic function in young male KO mice (Wooldridge et al. 2008). In these animals, SMTNL1 was also demonstrated to affect the transcriptional landscape of metabolic genes involved in glycolysis and glycogen storage (Lontay et al. 2015). Moreover, the deletion of SMTNL1 in young female mice promoted the switching of skeletal muscle from an oxidative to a glycolytic phenotype akin to what was observed in pregnant animals (Lontay et al. 2015). SMTNL1 can influence elements of thyroid hormone signaling, insulin signaling and glucose metabolism in C2C12 myotubes and potentially prevents hyperthyroidism-induced changes in skeletal muscle (Tamas et al. 2022; Major et al. 2021). However, the role of *Smtnl1* in the regulation of energy balance is largely unknown. Since SMTNL1 is most abundantly expressed in skeletal muscle (Wooldridge et al. 2008), and skeletal muscle plays a crucial role in systemic energy homeostasis (Baskin et al. 2015), we predicted that the global deletion of SMTNL1 might have an impact on the metabolic phenotype of ageing mice.

## MATERIALS and METHODS

### Animal Husbandry

Wild-type (WT, *Smtnl1*^+/+^) and knockout mice (KO, *Smtnl1*^-/-^, global) on a 129S6/SvEvTac background were employed (Wooldridge et al. 2008). Genotyping PCR was completed with primers (NeoPro: 5’-ACGCGTCACCTTAATATGC; *Smtnl1* WT: 5’- TTCACCTTTGACCC; and *Smtnl1* Rev: 5’-CAAAAGAGACCTGGC) and isolated genomic DNA from tail snips as previously described (Murali et al. 2023). Mice were housed in conventional shoe-box cages and provided a normal chow diet (5062 Pico-Vac® Mouse Diet 20, LabDiet, St. Louis, MO, USA) with a standard 12L:12D light-dark cycle within the Animal Health Unit at the University of Calgary. All animal use and protocols were approved by the Animal Care and Use Committee at the University of Calgary and conform to the guidelines set by the Canadian Council of Animal Care.

### Animals, housing and diets for metabolic phenotyping

Male and female *Smtnl1*^-/-^ mice (n = 8/group) at 12 months of age were housed individually in a Comprehensive Laboratory Animal Monitoring System (CLAMS®; Columbus Instruments, Columbus, OH, USA). The WT 129S6/SvEvTac background strain littermates served as controls (n = 8/group). Mice were provided a normal chow diet (5062 Pico-Vac® Mouse Diet 20; LabDiet, St. Louis, MO, USA) for at least 3 days during an initial acclimation period. The temperature (23–24 ℃) and humidity (21– 24 %) were controlled in animal rooms with a 12-h light/dark cycle. Food and water were provided *ad libitum*, except during times of fasting as noted below.

### Study design for metabolic phenotyping

In all cases, individual animals were subjected to continuous monitoring of energy intake, respiratory exchange ratio (RER), and energy expenditure(EE) with the CLAMS. After 3 days of acclimatization, the animals continued to be fed the normal chow diet (21% calories from fat, 55% calories from carbohydrate, and 24% calories from protein, 3.75 kcal/g; 5062 Pico-Vac® Mouse Diet 20; LabDiet, St. Louis, MO, USA) for 7 days. On day 5, animals were removed from the CLAMS and scanned using a Minispec LF-110 NMR Analyzer (Bruker Billerica, MA, USA) to determine body composition. To assess the basal metabolic rate of both genotypes, the mice were fasted for 24 h on day 5 prior to refeeding on day 6, with continuous monitoring of energy intake, EE, and RER. From day 8 to 15, the animals were provided a high fat diet that resembles a ‘Western diet’ (HFD: 40% calories from fat, 45% calories from carbohydrates, and 15% calories from protein, 4.63 kcal/g) for 8 days. To assess whether calorie depletion after *ad libitum* HFD feeding altered energy balance, the mice were not fed for 24 h on day 13 with subsequent return of HFD feed on day 14. The EE measured under the food- withdrawal condition was referred to as basal EE (Blais et al. 2018; Speakman 2013), and the expenditure following refeeding was the postprandial EE (Singh et al. 2019).

### Energy balance measurements

Food consumption, RER, and EE were monitored for 48 hours with CLAMS as previously described (Singh et al. 2016). Food spillage was manually recorded and used to correct the daily food (energy) intakes before completing analyses. The volume of oxygen consumed (VO2, ml/kg body weight/h), carbon dioxide produced (VCO2, ml/kg body weight/h), and RER, were recorded by indirect calorimetry (CLAMS; 2 L/min flow and ∼48 min sampling interval). The total EE was computed as previously published (Singh et al. 2018) using the following equation: VO2 × [3.815 + (1.232 × RER)].

### Body and liver composition

Body weight was recorded using a conventional laboratory scale. Whole body composition, comprising fat, lean and free fluid masses, was measured in unanesthetized mice at the end of low- and high fat diet feedings but before the 24-hour fasting period using a Minispec LF-110 NMR Analyzer (Bruker). The hepatic fat and lean mass were also determined with the NMR Analyser by scanning small portions (100 mg) of dissected liver tissue with the NMR probe. For every mouse/tissue sample, an average of 3 consecutive technical replicates were used.

### Intra-peritoneal glucose tolerance test

For the intraperitoneal glucose tolerance test (IPGTT), mice were fasted overnight (16 hours) at the end of day 15. An intraperitoneal injection of 50% (w/v) dextrose solution at a dose of 2 g/kg body weight was then administered, and blood glucose concentrations were determined from the tail vein using a handheld glucometer (Accu-Chek Glucose Meter; Roche, Basel, Switzerland). The area under the curve above the baseline (AUC) for blood glucose during IPGTT were calculated using the trapezoidal rule. The slope of the decline of plasma glucose values from the highest point during the IPGTT, calculated from the regression analysis provided an index of insulin sensitivity (Solberg et al. 2006).

### Statistical analyses

Repeated measures of calorie intake, RER, and EE were analyzed with linear mixed model and the appropriate covariance structures followed by Benjamini-Hochberg *post hoc* analyses using the Statistical Package for Social Sciences v.24.0 (IBM SPSS, Chicago, IL, USA). The model included the fixed effects of genotype and time, and their interactions where applicable. For EE, the sum of lean mass and 0.2 × fat mass was the covariate (Even and Nadkarni 2012). The difference between genotypes in body weight, whole body composition, tissue weights, liver tissue composition, and AUC were analyzed by unpaired Student’s t-test using GraphPad Prism v.9.4.0 (GraphPad Software, La Jolla, CA). Where appropriate, statistical assessments of more than 2 groups were completed by two-way ANOVA followed by Sidak’s multiple comparisons test. Data were expressed as means ± SEM (n = 5-9/group), and p < 0.05 was considered statistically significant.

## RESULTS

### Age-associated changes in body weight for male and female mice

Whole animal body weights were recorded from the age of 3 months. Consistent weight gain was observed for male mice, regardless of genotype (**Figure 1A**), until the age of 10 months. At this age, body weight stabilized. A general decline in body weight was observed for both male cohorts for ages >15 months. Although the male *Smtnl1*^-/-^ mice generally displayed higher body weights than their WT counterparts, the difference was significant only at 12 months of age. Females also displayed weight gain at young ages (**Figure 1B**); the body weight of WT mice plateaued after 9 months while the KO mice continued to gain weight until 15 months of age. Female *Smtnl1*^-/-^ mice displayed significantly greater body weights when compared to their WT littermates from the age of 11 to 16 months. After the age of 12 months, the body weight of both female cohorts displayed more variability. The body weights of female WT and KO mice displayed distinct age-related responses beyond 15 months of age; in this case, the female WT mice maintained a plateau at approximately 27 g while KO animals exhibited a steady decline back to WT values from a recorded peak weight of 34 g.

**Figure 1.**
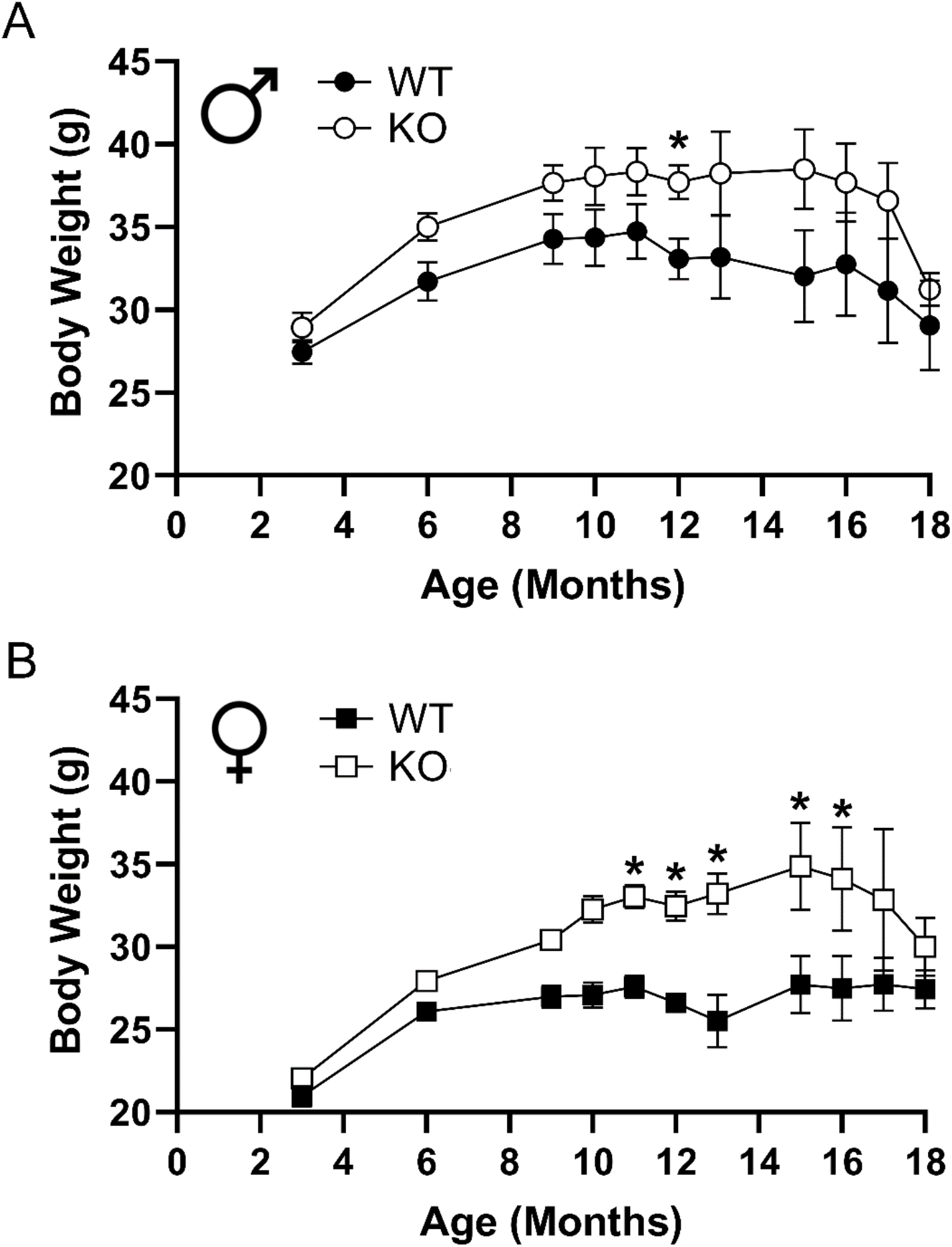
Body weights of wildtype and *Smtnl1^-/-^* mice with increasing age. Body weights of (**A**) male and (**B**) female wildtype (WT) and *Smtnl1^-/-^* (KO) mice from the age of 3 months to 18 months were recorded. Data are expressed as mean ± SEM of body weights measured monthly (n = 10-27/group) and analyzed using repeated measures 2-way ANOVA followed by Sidak’s multiple comparison test. The main effects- genotype, and time were significant (p < 0.0001) for both male and female cohorts but not their interaction. * p < 0.05 significant difference between genotypes at a given age.

### Whole body adiposity is enhanced with SMTNL1 deletion

At 12 months, of age, both male and female KO mice were significantly heavier than their WT litter mates. So, additional examinations of body composition were completed for these animals using a non-invasive, full body NMR scanner. In males, no difference was observed between genotypes with respect to percent fat and lean mass (**Figure 2A**). Interestingly, female KO mice had significantly increased percent fat mass (2-fold) with a concomitant decrease in both percent lean and fluid masses (**Figure 2B**).

**Figure 2.**
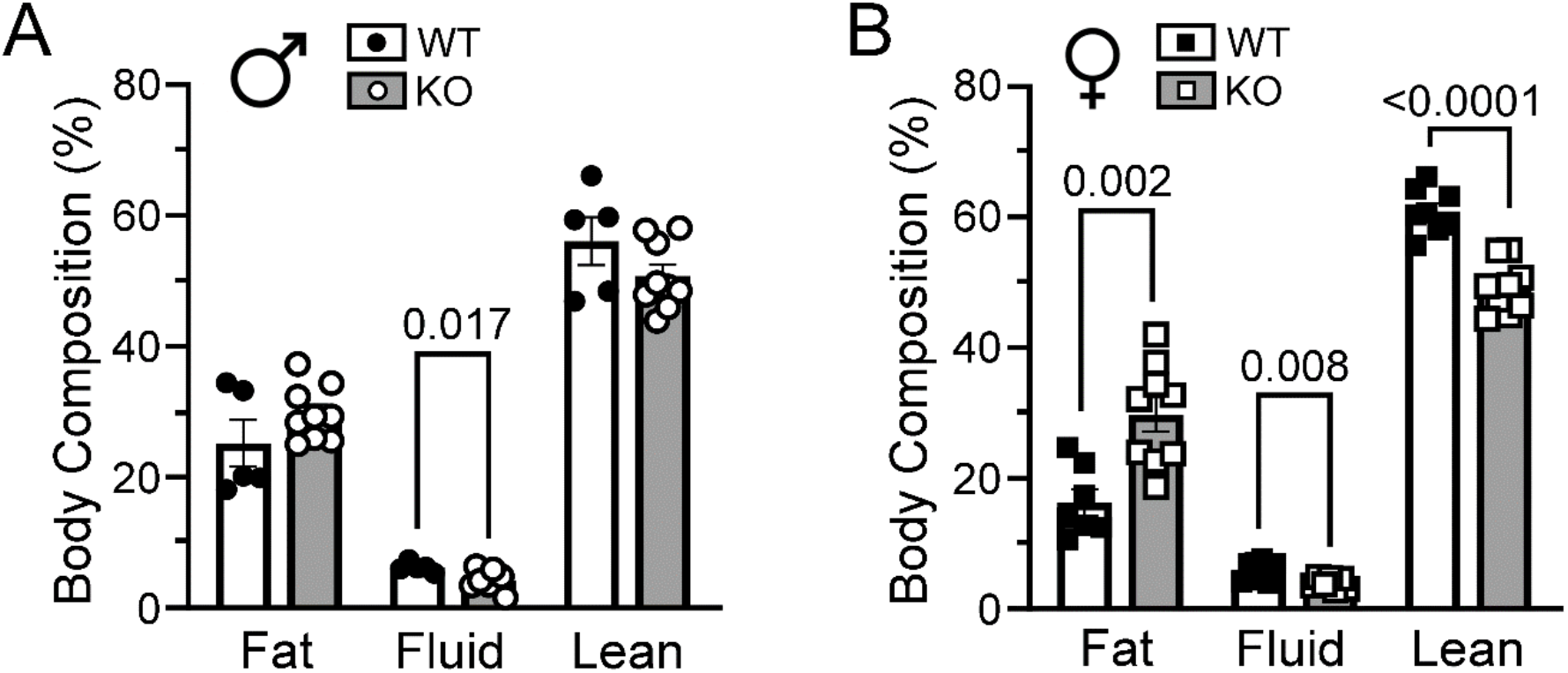
Effect of SMTNL1 deletion on body composition of ageing mice. The body composition of 12-month-old animals was measured using a Minispec LF-110 NMR Analyzer. Both wildtype (WT) and *Smtnl1^-/-^* (KO) mice were provided a normal chow diet. Fat and lean mass as % of body weight is given for male (**A**) and female (**B**) mice. Data are expressed as mean ± SEM (n = 5-9/group) and analysed using unpaired Student’s t-test. The numbers above the bars indicate the p values with p < 0.05 considered significant.

### Enhanced energy intake in male and female Smtnl1^-/-^ mice fed a high fat diet

Food consumption was recorded for male and female cohorts (WT and KO, 12 months of age) and used as a surrogate for energy intake. A two-way ANOVA with repeated measures revealed significant effects of genotype (p<0.0001) on the cumulative energy intake for males fed a normal chow, but not for females (p=0.941). In support of the main genotype effects, male mice of both genotypes steadily consumed normal chow during the refeeding period after fasting (**Figure 3A,i**); food consumption continued even after conclusion of the dark period, which is the active feeding time for rodents. Similarly, the female mice, irrespective of the genotype, displayed comparable increases in energy intake post-fasting, during the dark period and beyond in the normal chow diet group (**Figure 3A,ii**). When provided a high fat diet, the main effects (i.e., genotype and time) were significant for the cumulative energy intake for males (**Figure 3B,i**) and females (**Figure 3B,ii**). However, the hyperphagic response to refeeding was greater in the KO mice of both sexes when compared to WT littermates.

**Figure 3.**
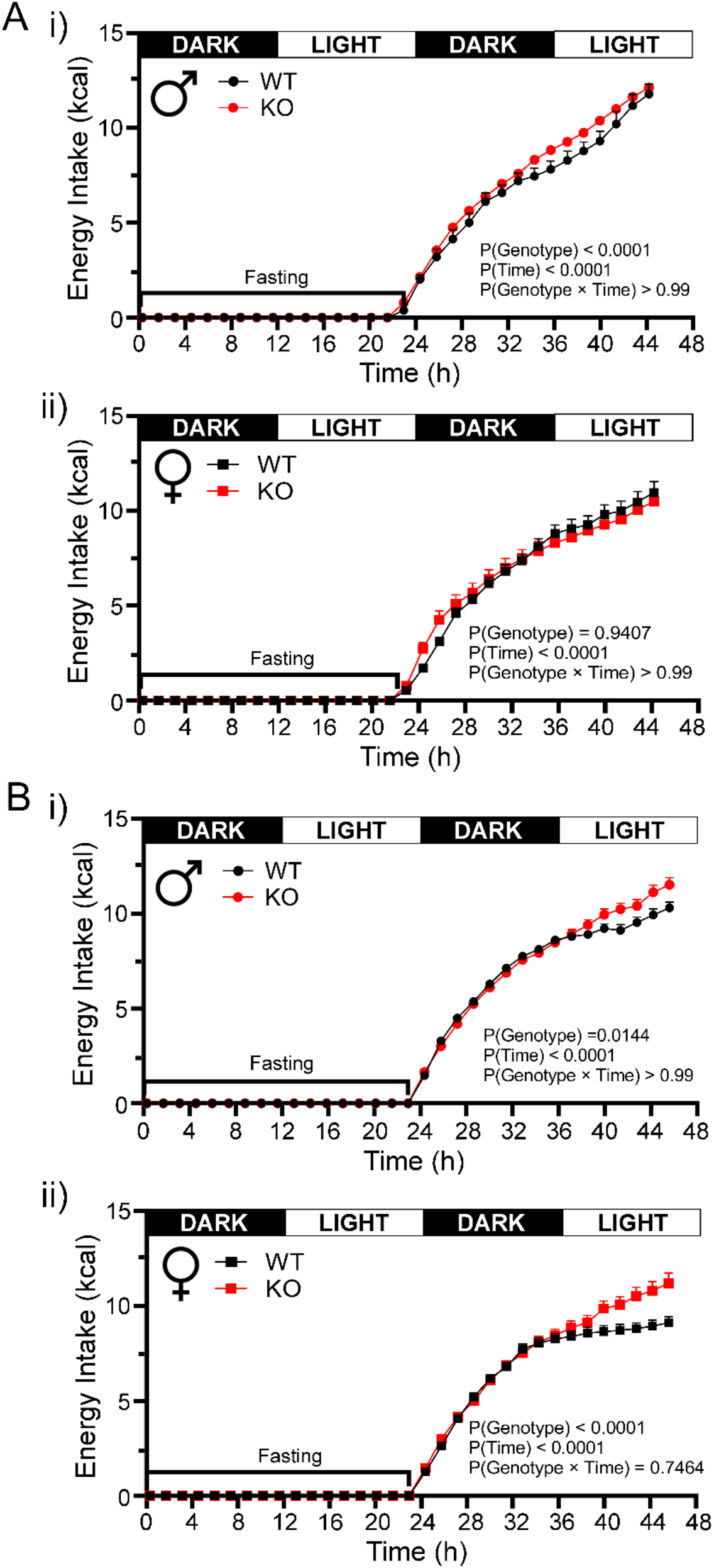
Effect of SMTNL1 deletion on energy intake of 12 month old male and female mice. Wildtype (WT) and global *Smtnl1^-/-^*(KO) mice were acclimated within individual CLAMS chambers with *ad libitum* provision of normal chow. On day 5, the mice were fasted for 22 hours and refed with normal chow with measurement of energy intake (**A**: *panel i*, males; *panel ii*, females). Thereafter, the mice were provided a high fat diet for 8 days. On day 13, the mice were again fasted for 22 hours and refed a high fat diet (**B**: *panel i*, males; *panel ii*, females) with subsequent recording of energy intake. Data are expressed as mean ± SEM (n = 5-9/group) and analyzed using linear mixed models for repeated measures followed by Benjamini-Hochberg *post hoc* analyses . All the main effects and interactions were considered significant with p < 0.05.

### Increased energy expenditure of male Smtnl1^-/-^ mice fed a normal chow diet and female Smtnl1^-/-^ mice fed a high fat diet

Energy expenditure (EE) was measured with indirect calorimetry of unrestrained mice. With normal chow provision, the main effects of genotype and time on basal EE and postprandial EE and their interactions were significant for the male study group (p < 0.0001). The basal EE monitored during the fasting period was higher in the KO mice than in the WT group; this response was reversed in the postprandial dark period where the WT mice had higher EE than the KO mice (**Figure 4A,i**). A similar response pattern was observed for male animals provided with a high fat diet where the basal EE was higher in the absence of SMTNL1, and the postprandial EE was higher in the WT genotype during the dark period (**Figure 4A,ii**). Ultimately, the interaction between genotype and time for high fat feeding was significant only for the postprandial EE suggesting that the effect of time on the EE differed between the genotypes only during refeeding and not during fasting. On the other hand, the main effects of genotype and time on basal EE and postprandial EE and their interaction were significant for female groups fed either normal (**Figure 4B,i**) or high-fat (**Figure 4B,ii**) diets. Interestingly, both basal and postprandial EE were higher for the female KO animals provided normal chow when compared with the WT littermates (**Figure 4B,i**). This difference between female cohorts was further exaggerated during high fat refeeding in both the dark and light period (**Figure 4B,ii**).

**Figure 4.**
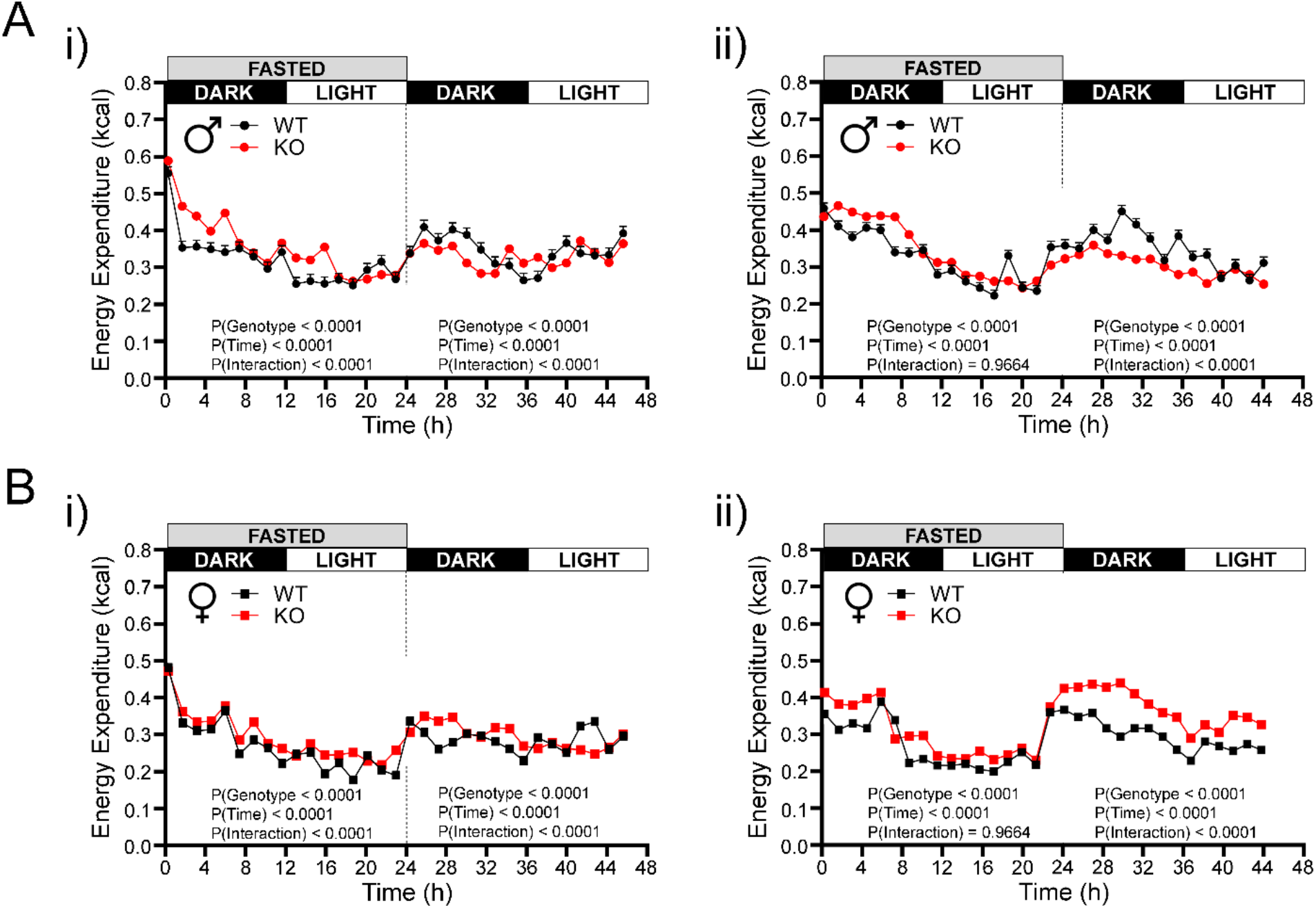
Effect of SMTNL1 deletion on energy expenditure of 12 month old male and female mice fed normal chow and high fat diets. Energy expenditure was monitored for 12-month-old, wildtype (WT) and *Smtnl1^-/-^* (KO) mice over 48 hours. Male (**A**) and female (**B**) mice were provided normal chow (*panel i*) or high fat (*panel ii*) diets and the monitoring period was initiated with 22 hours of fasting. Data are expressed as mean ± SEM (n = 5-9/group) and analyzed using linear mixed models for repeated measures followed by Benjamini-Hochberg post hoc analyses separately during and after the fasting period. All the main effects and interactions were considered significant with p < 0.05.

### Altered respiratory exchange ratio of male and female Smtnl1^-/-^ mice fed low and high fat diets

The respiratory exchange ratio (RER) was monitored as an indicator of metabolic fuel preference. For the male mice provided normal chow feeding (**Figure 5A,i**), both WT and KO cohorts displayed a decrease in RER from ∼ 0.9 to 0.75 in the light period during fasting that was indicative of a switch from carbohydrate to mixed fat use. RER returned to ∼ 0.9 during the refeeding phase for both genotypes in both dark and light periods. The repeated measures statistical analysis showed significance only in the main effects of time. In the male cohorts provided a high fat diet (**Figure 5A,ii**), RER was reduced after the dark period during the fasting phase to levels suggestive of mixed fat use, but this reduction was more apparent for the WT mice than the KO mice. In this case, the RER of male KO mice did not drop below a value of 0.8 whereas the corresponding RER of WT mice gradually declined to ∼ 0.7, suggesting that KOs prefer to utilize carbohydrates when fed high fat diets. The repeated measures statistical analysis showed the main effects of genotype and time were significant but not their interaction. During the refeeding phase, RER increased substantially with KO mice rebounding to higher RER (i.e., ∼1.0) than the WT counterparts. There was a noticeable decline in RER for both WT and KO cohorts during refeeding in the light phase; however, the values for male KO mice remained elevated above those of the WTs (**Figure 5A,ii**).

**Figure 5.**
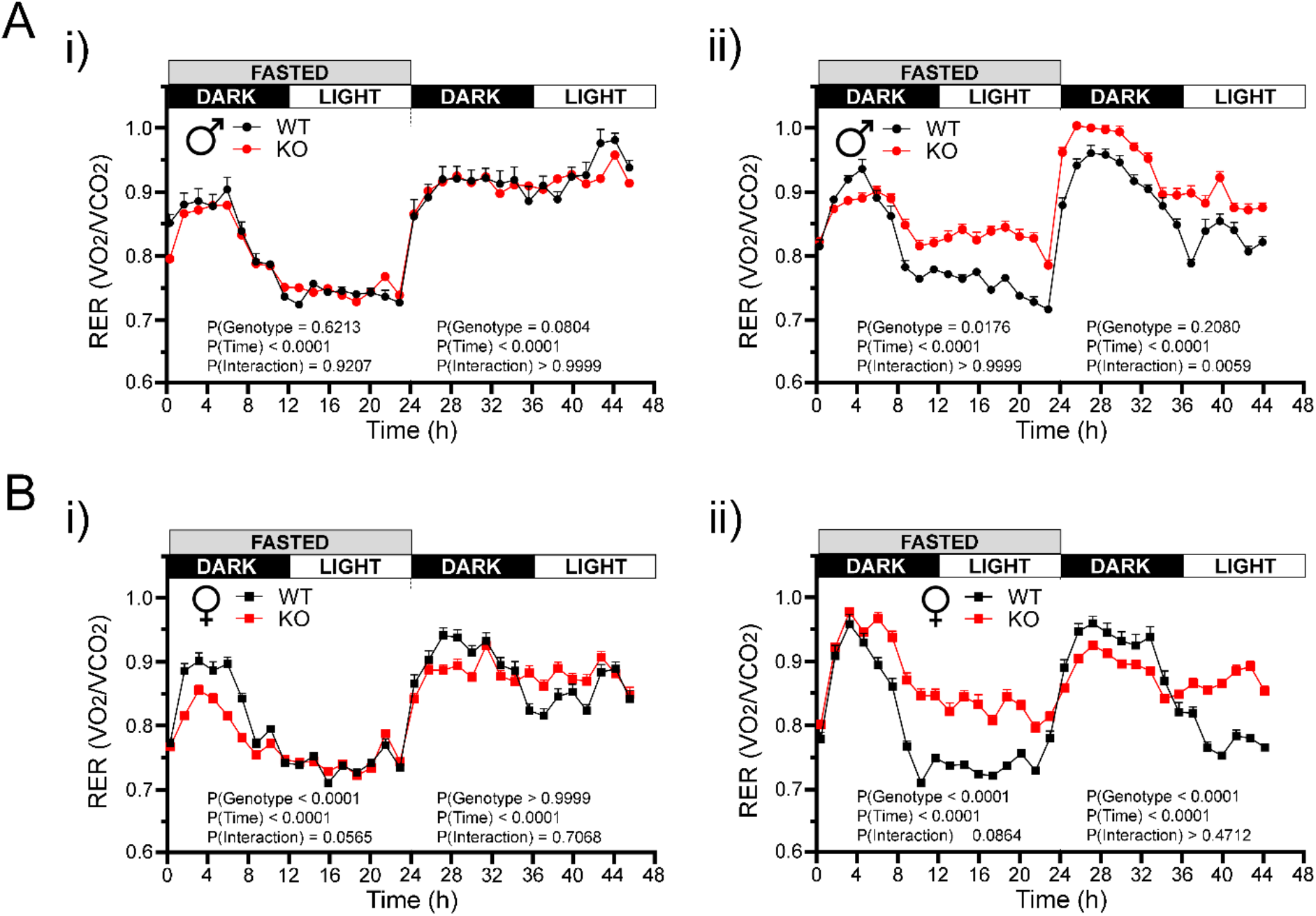
Effect of SMTNL1 deletion on respiratory exchange ratio of 12 month old male and female mice fed normal chow and high fat diets. Following three days of acclimation, the respiratory exchange ratio (RER) was monitored for 12-month-old, wildtype (WT) and *Smtnl1^-/-^* (KO) mice over 48 hours. Male (**A**) and female (**B**) mice were provided normal chow (*panel i*) or high fat (*panel ii*) diets and the monitoring period was initiated with 22 hours of fasting. Data are expressed as mean ± SEM (n = 5-9/group) and analyzed using linear mixed models for repeated measures followed by Benjamini-Hochberg post hoc analyses separately during and after the fasting period. All the main effects and interactions were considered significant with p < 0.05.

The female cohorts fed a normal chow diet generally displayed similar metabolic responses to fasting and refeeding in dark and light phases as their male counterparts (**Figure 5B,i**). The RER of both genotypes similarly decreased in the fasted light period to suggest mixed fat use. However, the RER of female WT mice in the initial fasted dark period was higher than that of the KO cohort. The repeated measures statistical analysis showed the significant main effects on the RER were genotype and time (p < 0.0001). During refeeding, the RER of the female WT cohort was elevated in the dark period followed by a reduction in the light period. In addition, the female KO mice provided normal chow displayed a unique plateau of RER at ∼0.9 throughout the refeeding phase. In contrast, the RER of female WT mice dropped from ∼0.95 to be near 0.8 during the light phase. In the female cohorts provided a high fat diet (**Figure 5B,ii**), marked differences were apparent between genotypes in the fasted light phase, with KO mice showing markedly higher RER than the WT mice that support an increased utilization of carbohydrates over fat as fuel in the absence of SMTNL1. During refeeding, the RER of the WT cohort rebounded significantly in the dark phase, from RER values of 0.75 to 0.95. The RER was higher than the KO mice, but this pattern was reversed in the subsequent light period with a decline in RER only found for the WT cohort. The statistical analysis revealed the main effects of time and genotype on RER to be significant but not their interaction.

### White adipose tissue weights are increased in male and female Smtnl1^-/-^ mice

The total weight of white adipose tissues (WAT) dissected from KO mice was substantially increased for both male and female cohorts (**Figure 6A,B**). Although total WAT content was significantly elevated with Smntl1 knockout in both sexes, the female mice displayed a larger distinction in WAT content between genotypes (**Figure 6B**). In this case, the relative WAT mass increased from 6 to 16% of body weight for WT and KO mice, respectively. The final wet weight of the liver was also measured, but no differences between genotypes were noted for either sex when values were normalized to body weight (**Figure 6A,B**). The findings of increased adiposity were aligned with the non-invasive body composition data collected with NMR imaging (**Figure 2**). Among the individual fat pads dissected from the male KO mice, all except the epicardial and brown adipose tissues (BAT) were increased when compared to the WT littermates (**Figure 6C**). These genotypic differences in individual fat pad weights were also recorded in the female sex with an increase in epicardial adipose tissue also observed (**Figure 6D**). The female-specific parametrial fat depot, the largest visceral fat pad in female mice and the closest analog to human omental adipose tissue, was elevated by 3-fold in the KO animals. The male-specific epidydimal fat depot was also enlarged with deletion of *Smtnl1*.

**Figure 6.**
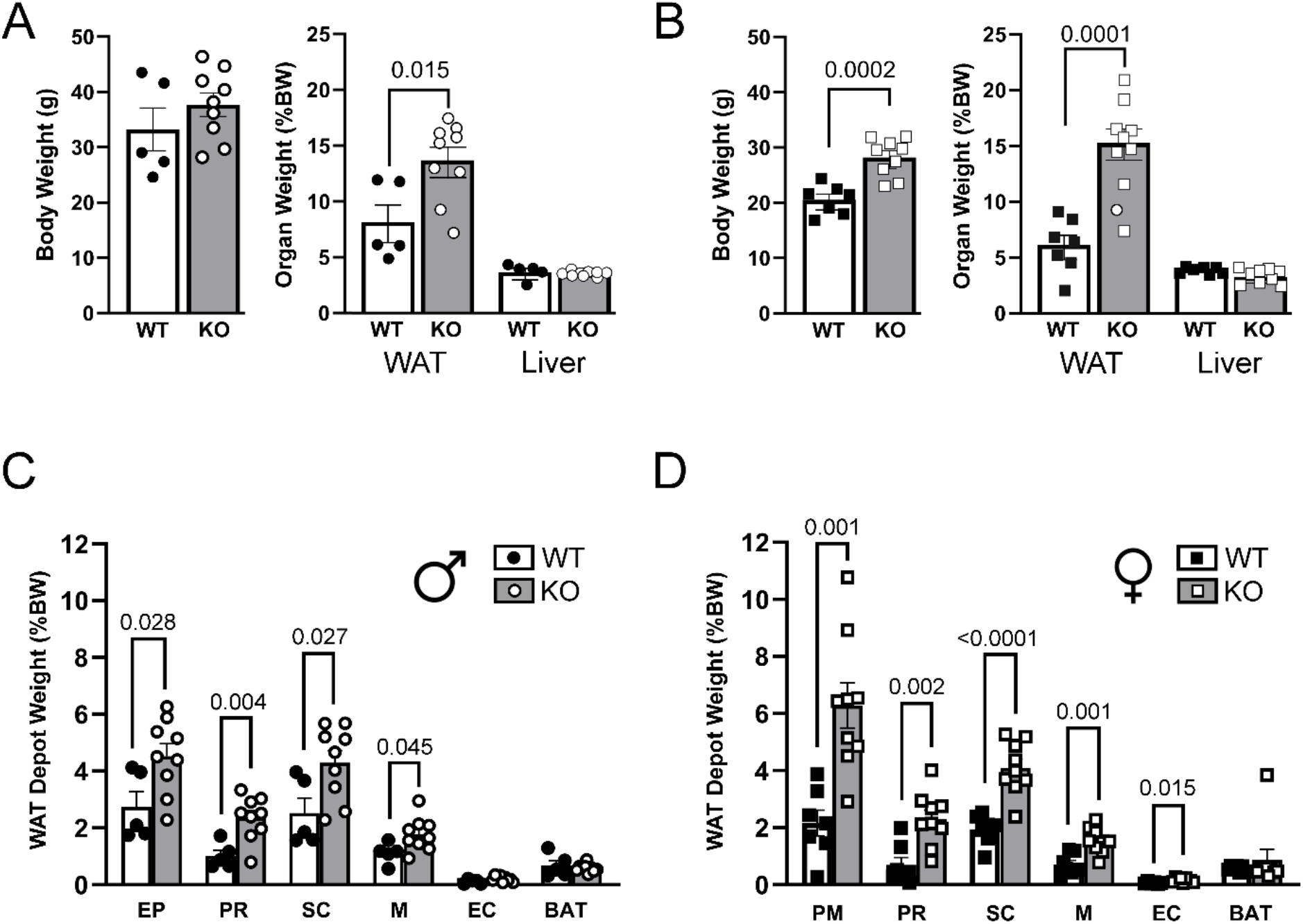
Comparison of body weights and tissue adiposity between genotypes of aged male and female mice. The final body weight was measured for male (**A**) and female (**B**) mice after measurements of energy balance and body composition parameters with CLAMS for wildtype (WT) and *Smtnl1^-/-^* (KO) animals. Liver and total white adipose fat pad (WAT) weight were also compared between the genotypes of both sexes. The weights of individual fat pads were determined for both genotypes of male (**C**) and female (**D**) mice: including, epididymal (EP, in males only), peri-renal (PR), subcutaneous (SC), mesenteric (M), epicardial (EC), brown adipose (BAT), and parametrial (PR, in females only). Data are expressed as mean ± SEM (n = 5-9/group) and analyzed using an unpaired Student’s t-test. The numbers above the bars signify the p value where p < 0.05 was considered significant.

### Delayed clearance of postprandial blood glucose in the absence of SMTNL1 in female mice

Intraperitoneal glucose tolerance (IPGTT) was used to examine the impact of SMTNL1 deletion on the glucose metabolism of ageing male and female mice. For the 12 month old male mice, IPGTT revealed no difference in glucose clearance between genotypes (**Figure 7A,i**); a similar increase in blood sugar after 30 minutes of bolus injection was observed. Notably, the baseline blood glucose level of the KO mice was lower than the WT mice (**Figure 7A,ii**; 5.84 ± 0.19 mmol/L vs 4.30 ± 0.30 mmol/L, p < 0.005); however, no distinctions in the slope of the decay response for plasma glucose during IPGTT (**Figure 7A,i**) or the area under the curve (AUC) above baseline (**Figure 7A,iii**) were noted. In the female cohort, KO mice showed signs of glucose intolerance with substantially elevated blood sugar levels at 30 minutes after the glucose bolus when compared to the WT counterparts (**Figure 7B,i**). The significant interaction between time and genotype confirmed that the effect of time on the levels of blood sugar differed between animals of different genotypes in the female cohort. The female SMTNL1 KO mice also displayed significantly lower baseline blood glucose levels at the start of the testing (**Figure 7B,ii**; 5.79 ± 0.30 mmol/L vs 3.71 ± 0.11 mmol/L, p < 0.001). The AUC above baseline was not different between the female WT and SMTNL1 KO mice (**Figure 7B,iii**); however, the slope of the decline of blood glucose levels from the highest point, an index of insulin sensitivity was almost doubled in the KO animals (**Figure 7B,i**).

**Figure 7.**
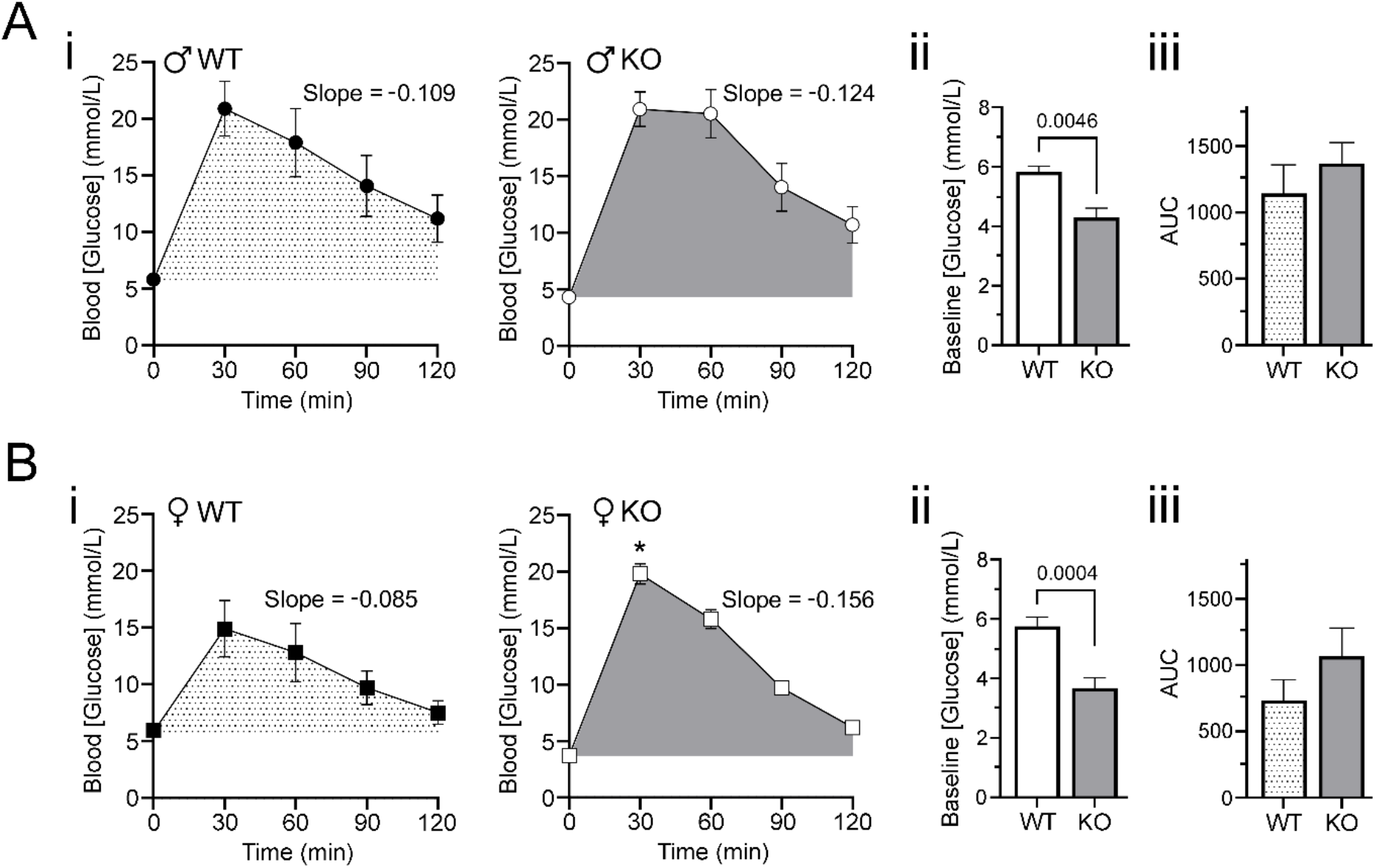
Comparison of glucose tolerance between ageing *Smtnl1^-/-^* and wildtype male mice. Intraperitoneal glucose tolerance was tested at 12 months of age for wildtype (WT) and *Smtnl1^-/-^*(KO) mice. Male (**A**) and female (**B**) animals were fasted overnight for 16 hours, and blood glucose levels were measured 30, 60, 90 and 120 min after administration of a glucose bolus (time 0; pre-glucose bolus baseline level). The baseline glucose concentration (**A** & **B**; *panel ii*), area under the curve above the baseline (AUC: **A** & **B**; *panel iii*) were calculated and analyzed with an unpaired Student’s t-test (p < 0.05 was considered significant). The slope of the decline of blood glucose levels from the 30 min timepoint was calculated using regression analysis, as an index of insulin sensitivity. Data are expressed as mean ± SEM (n = 5-9/group).

## DISCUSSION

We provide key evidence in support of a sexually dimorphic role for SMTNL1 in modulating energy balance. First, SMTNL1 KO mice had greater weight, adipose mass, and glucose intolerance, which is particularly prominent in females and exacerbated by high-fat feeding. Second, the higher adiposity in the SMTNL1 KO female mice is likely driven by a greater hyperphagic response that is unmasked on a highly palatable high-fat diet and a coincident suppression of basal energy expenditure. In males, despite marginally heavier weights and postprandial hyperphagia, the SMTNL1 KO mice had greater basal energy expenditure that might have limited adipose accretion. Together, these findings support the notion that endogenous SMTNL1 is important for promoting satiety and thermogenesis, particularly in females.

The evident gain in body weight in the older, 12 month females is suggestive of increased adiposity with knockout of SMTNL1. This genotypic difference in body composition was even more drastic upon acute high fat diet feeding. Unlike their female counterparts, male KO mice had only a slight increase in fat mass upon high fat diet feeding, which was not significant. To evaluate if the advancing age was a factor in the influence of SMTNL1, additional body composition analyses were completed with young mice (i.e., aged 3 months). Like the body weight data, the whole-body lean and fat contents were not different between genotypes in the young male cohort (**Figure 8**). In addition, the % fat content of the KO females was not significantly different from the WT and KO males although lean body mass was decreased. Interestingly, the 12-month age point in female mice is suggested to correspond with the menopausal human age (Diaz Brinton 2012), and so, the observation is in line with the fact that women tend to gain weight and fat mass around the menopausal age with a decline in systemic estrogen levels (Simkin-Silverman and Wing 2000). This is further supported by studies on 18-month-old aged male mice, which demonstrated significant reduction in body mass, visceral adiposity and ectopic lipid deposition upon estrogen supplementation (Stout et al. 2017).

**Figure 8.**
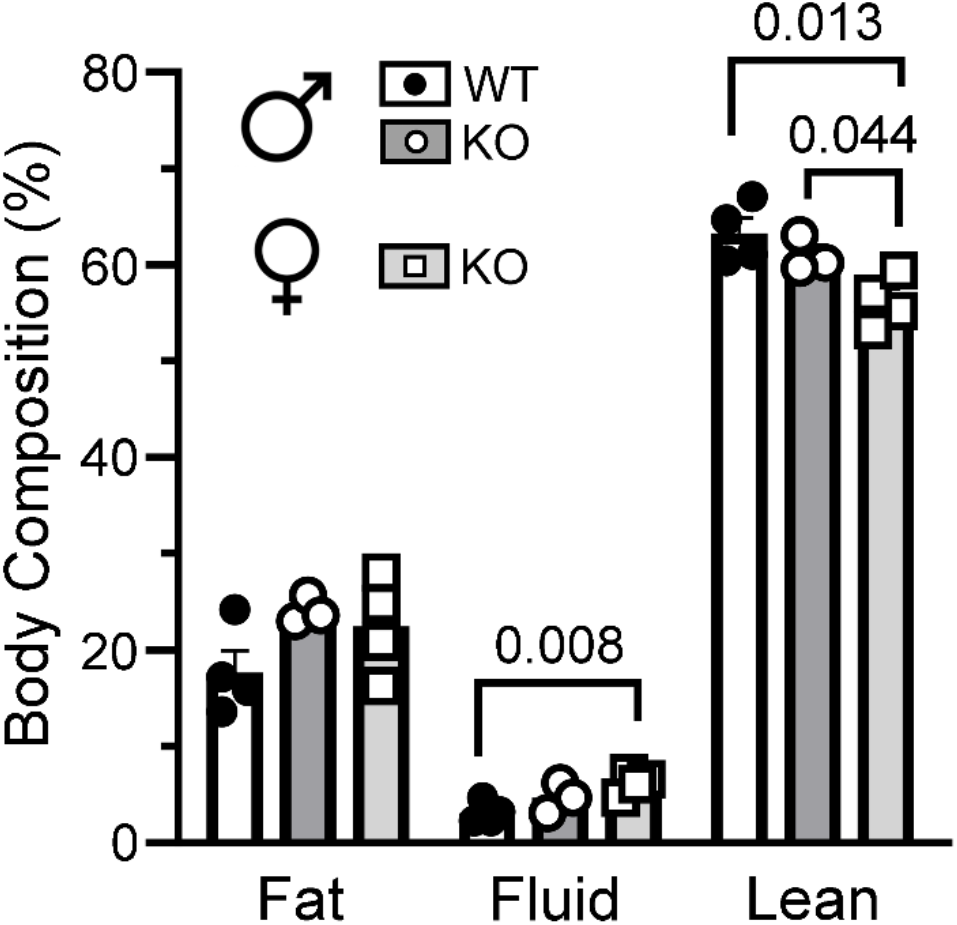
Comparison of fat mass and lean mass between young male and female *Smtnl1^-/-^* mice. The body composition of 3-month-old mice was measured using Minispec LF-110 NMR Analyzer. Fat, fluid and lean mass as % of body weight is given for male and female mice. Data are expressed as mean ± SEM (n=3-5/group) and analysed using one-way ANOVA followed by Tukey’s multiple comparison test. The numbers above the bars show the p values where p < 0.05 was considered significant.

Metabolic maladaptation often leads to elevated adiposity, and additional investigations examined the impact of SMTNL1 deletion on energy intake, energy expenditure and substrate utilization in these middle-aged animals. Intriguingly, the cumulative food intake of male mice which rose sharply during the dark period (i.e., the normal activity and feeding period) did not plateau in the subsequent light period in both normal chow diet and high fat diet fed groups. In the female cohort, there was no difference between KO and WT in the energy intake, both of which steadily increased beyond the active dark period when fed a normal chow diet. This was similar to results obtained previously for a young female cohort of SMTNL1 KO mice, where no genotypic difference was observed for food intake while on a normal chow diet (Lontay et al. 2015). In the current study that provided mice with a high fat diet, the food intake of the WT animals plateaued after the dark period while the KO animals continued to progress. Thus, the deletion of SMTNL1 seemed to delay satiety in mice fed a high fat diet, which was more pronounced in the female mice. The present observation of elevated energy intake could partially explain the higher adiposity observed in the female KO mice. Indeed, caloric restriction is a proven strategy to reduce visceral adiposity and thereby alleviate insulin resistance in aged rodents (Barzilai et al. 1998). Whether the deletion of SMTNL1 affects any of the adipokine levels such as leptin that promote the sensation of satiety, needs to be further investigated. Circulatory levels of leptin have been, however, shown to be unaffected by diet in middle-aged mice (Duncan et al. 2016) although it is increased with age in men (Van Den Saffele et al. 1999).

Another metabolic parameter energy expenditure (EE), under basal condition in males (i.e., during fasted dark phase) was markedly higher in the absence of SMTNL1 and this was reversed with refeeding where WTs had higher post-prandial EE in the active dark phase. This observation was more pronounced when males were fed a high fat diet. However, it is to be noted that elevated energy intake in the light period of high fat diet fed males was not accompanied by increased EE. The basal and postprandial EE in females were slightly higher in the KO group and became markedly high upon high fat diet feeding. The overall enhanced EE in heavier SMTNL1 KO animals is not surprising as obesity is shown to be linked with an elevated daily EE and increased basal metabolic rate in proportion to the excess body weight (Balasse 1994; Careau et al. 2021). Additionally, excess body weight was proved not an outcome of reduced sleeping, resting EE, and fat metabolism (Weinsier et al. 2003). This further justified hyperphagia or higher energy intake of the KO mice in order to maintain a weight-gain state while the daily energy expenditure was greater than their WT counterparts.

In normal physiology, increases in RER toward 1.0 is indicative of a shift towards carbohydrate usage and decreased RER means increased use of fatty acids. The current findings of reduced RER during the fasting phase agreed with literature reports where increased RER reflects primary carbohydrate utilization with feeding and decreased RER indicates increased body fat metabolism during fasting in metabolically flexible subjects (Storlien et al. 2004). Earlier work revealed a hypermetabolic shift towards carbohydrate utilization for young SMTNL1 KO female mice in the fed state (Lontay et al. 2015). Similarly, in the present study, the middle-aged female KO mice on high fat diet had higher RER, suggesting a significant shift to utilization of carbohydrates as fuel during the fasted diurnal cycle and the fed light period. When fed a normal chow diet, KO mice irrespective of the sex, had reduced RER during the diurnal cycle when compared to WT littermates fed a high fat diet. The findings support the ability of KO mice to effectively shift between substrates when on a normal chow diet; however, upon high fat feeding the KO animals become metabolically less efficient and display a preference for carbohydrate catabolism. Generally, HFD challenges the oxidative machinery of the skeletal muscle and metabolically efficient subjects adapt by increasing fatty acid oxidation (Ukropcova et al. 2005). Further, there was evidence of sex dimorphism in metabolic efficiency in middle-aged animals since the female WTs had lower RER than their male counterparts. Although no gene expression studies were conducted in the current investigation, a previous report using young WT and KO mice linked the deletion of SMTNL1 to the altered expression of genes associated with glycolysis, gluconeogenesis, glycogenolysis, glycogenesis, as well as fatty acid synthesis and oxidation in the skeletal muscle (Lontay et al. 2015). Similar effects on gene expression were also identified in the uterine smooth muscle type through its interaction with progesterone receptor (Bodoor et al. 2011).

A recent report has profiled morphometric and metabolic phenotypes of male C57BL/6N mice ranging from 3 to 36 months of age (Petr et al. 2021). Although female mice were not profiled, the body composition measurements and *in vivo* metabolic function of the aging male mice are in general agreement with the WT cohort used in this study. For example, % lean body mass and the ratio of subcutaneous/visceral fat tended to be higher with aging in the C56BL/N6 mice. Moreover, an age-dependent reduction in many parameters (e.g., blood glucose, body weight, % fat) was observed at ∼19 months of age. Another study of aging C57BL/6J mice provided by Chaix and co-authors (Chaix et al. 2021) suggests that the lean mass as a percentage of body weight was similar in male and female mice within the same age group; however, older mice of both sexes had reduced lean mass compared to younger mice (12-month vs. 3 month mice, respectively).

SMTNL1 deletion is associated with switching of skeletal muscle fibre-type from an oxidative to a glycolytic isoform in young female mice (Lontay et al. 2015). If this phenomenon could be extrapolated to middle-aged mice, it explains the increased shift towards carbohydrate utilization for fuel observed in this study. A preference for carbohydrate substrate would be predicted to result in reduced fatty acid oxidation and increased storage of excess free fatty acids as triglycerides in adipose tissue. In addition, the white adipose tissue of mice can further enlarge and become insulin resistant in the presence of excess calories in the form of dietary lipids and carbohydrates. This is in alignment with the present observation of increased body weight, whole body fat mass content and the total fat pad weights of SMTNL1 KO mice. However, the distribution of white adipose tissue depots rather than total adiposity is critical in predicting the cardiometabolic health risk (Goossens 2017). Hence, individual white and brown adipose tissue depots were examined revealing enhanced adiposity in visceral as well as subcutaneous adipose tissue depots in both male and female animals with SMTNL1 deletion. Specifically, the expansion of visceral adipose tissue was more prominent than the subcutaneous fat pad, with this difference being more explicit in the female KO mice. It is to be noted that the proximity of visceral fat to liver and heart facilitates the direct flow of various secretory molecules such as cytokines, adipokines and free fatty acids from the adipose tissue to these vital organs and can critically affect its function than the subcutaneous fat depots (Nielsen et al. 2004).

The excessive enlargement of WAT can adversely affect its secretary function (Huffman and Barzilai 2009; Sun, Kusminski, and Scherer 2011). In this case, the tissue phenotype becomes pro-inflammatory and insulin resistant; characteristics that are associated with the downregulation of glucose uptake and *de novo* lipogenesis and the ultimate development of hyperglycemia and glucose intolerance. This was apparent for the female SMTNL1 KO animals that show delayed clearance of postprandial blood glucose, overt metabolic imbalance and excessive adiposity. Impaired glucose tolerance was also previously reported in young female mice in the absence of SMTNL1 (Lontay et al. 2015). In contrast, the fasting blood glucose levels were suppressed in both sexes of SMTNL1 KO mice when compared to their WT counterparts.

While one-third of post-prandial glucose is absorbed by liver, the remainder is taken up by peripheral tissues, predominantly skeletal muscle via an insulin-dependent mechanism (Baron et al. 1988). When the dietary intake of high energy macronutrients exceeds the muscle glucose uptake and fatty acid oxidation capacity, insulin resistance and glucose intolerance follow. Moreover, the anti-lipolytic effect of insulin is also impaired. The subsequent hyperlipidemic state is driven by increasing lipolysis in adipocytes and the enhanced free fatty acid release into the circulation. Concurrently, insulin-sensitive hepatocytes continue to take up blood sugar and undergo *de novo* lipogenesis, finally leading to hepatic insulin resistance and eventually to non-alcoholic fatty liver disease (Hudgins et al. 2011). In the present study, even though SMTNL1 deletion was associated with a significant reduction in normalised liver weight in 12 month old male mice (**Figure 9A**), the hepatic % fat remained unchanged when compared to age-matched WT mice. The liver mass of 12 month old female KO mice was also reduced, similarly to that of males. Only the young female KO animals showed a significant increase, albeit modest, in the hepatic fat content (**Figures 9B**). Thus, it does not appear that the metabolic alterations and adiposity observed in older female *Smtnl1*^-/-^ mice are reflective of fatty liver.

**Figure 9.**
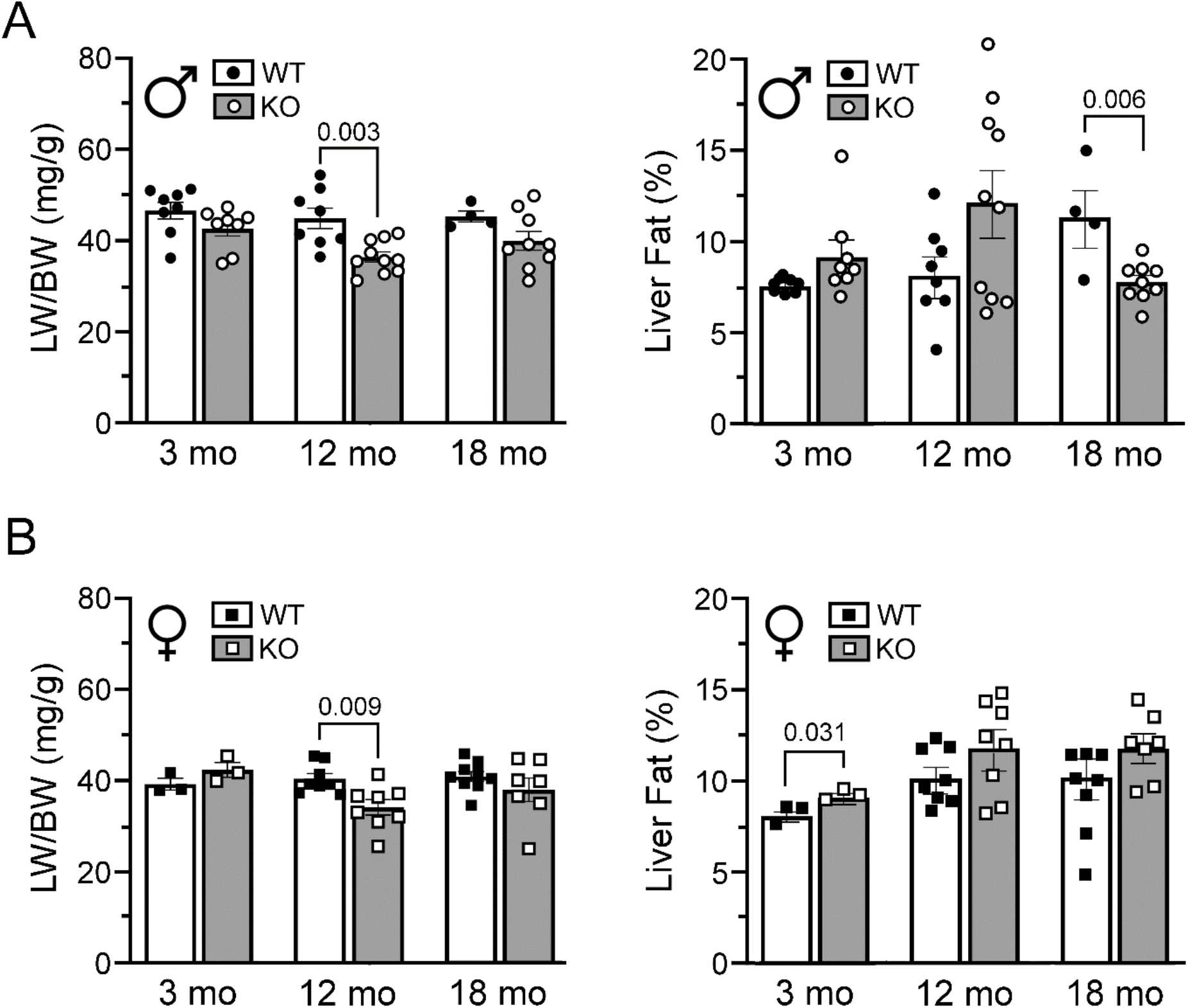
Comparison of hepatic weight and fat content between *Smtnl1^-/-^* and wild type mice with increasing age. Liver weight (LW) and % liver fat were determined for 3 different age groups of male (**A**) and female (**B**) mice: 3-month-old, 12-month-old, and 18-month. After gross dissection of livers, the liver weight (LW) was normalised to body weight (BW). Hepatic tissue samples (∼ 100 mg) were collected by punch biopsy. The tissue samples were scanned for fat mass composition using a Minispec LF-110 NMR Analyzer. Data are expressed as mean ± SEM (n = 4-10/group) and analysed using an unpaired Student’s t-test. The numbers above the bars signify the p value where p < 0.05 was considered significant.

A recent *in vitro* study reported an insulin-sensitizing role for SMTNL1 with its overexpression in C2C12 myotubes, mainly by the activation of Akt and induction of glucose transporter (Glut-4) expression (Tamas et al. 2022). The overexpression study showed that in the presence of progesterone, SMTNL1 could downregulate novel PKCε as well as JNK1 and relieve the inhibitory phosphorylation of insulin receptor substrate 1 at Ser318 and Ser307. Therefore, the removal of SMTNL1 could result in insulin resistance, leading to metabolic imbalance and adipose accumulation. Furthermore, inflammation associated with obesity is known to activate the JNK pathway and cause insulin resistance in the peripheral tissues by blocking downstream insulin signaling (Aguirre et al. 2000).

There was evidence that young female SMTNL1 KO mice were metabolically less efficient as they preferentially utilized carbohydrates as fuel and demonstrated impaired glucose tolerance (Lontay et al. 2015). These findings are now extended to older, 12 month female mice and demonstrate that the knockout of SMTNL1 leads to progressive morphometric alterations during aging (i.e., increased body weight, whole body fat mass, and altered cutaneous/visceral white adipose tissue ratios). These events were associated with hyperphagia, enhanced energy expenditure in the diurnal cycle and a greater shift towards carbohydrate utilization as energy substrate, rendering the KO animals metabolically less efficient upon high fat feeding. Taken together, the current findings establish a novel role for SMTNL1 in modulating adiposity and energy metabolism with ageing in a sex dimorphic way.

Some limitations of the present study were that the duration of the high fat diet intervention was acute (< 1 week) and that the same animals were sequentially reused within each dietary experiment. Nonetheless, clinical studies have followed similar strategies where after 1-5 days on control or high-carbohydrate diet, dietary fat was increased for 3-4 days in healthy, young men and women, and a positive fat balance was reported because of an imbalance between fat intake and fatty acid oxidation (Galgani et al. 2010; Schrauwen et al. 2000). A parallel evaluation of the metabolism of ageing male and female C57BL/6J mice supports distinct sex- and age-dependent responses to 3 months of Western diet feeding (Chaix et al. 2021). Namely, female mice displayed reduced insulin levels and elevated adipose tissue inflammation in response to Western diet feeding. Intriguingly, caloric restriction (i.e., time-restricted feeding) could protect both sexes against fatty liver and glucose intolerance while body weight benefits were observed only for males, irrespective of age. In additional, EE was also significantly affected by the feeding paradigm in male mice but not in females, irrespective of age. Whether provision of long-term

Western HFD high fat diet to SMTNL1 KO mice causes more overt metabolic syndrome with potential amelioration by calorie restriction or estrogen supplementation in ageing animals with SMTNL1 deletion remains to be investigated.

## ACKNOWLEDGEMENTS

The authors appreciate the assistance of Mona Chappellez in the monitoring and management of mouse breeding colonies.

## FUNDING STATEMENT

M.M. was supported by an Achievers in Medicine Doctoral Scholarship from the University of Calgary. This work was supported by research grants from the Canadian Institutes of Health Research (MOP#97931 to J.A.M.) and the American Heart Association (#953881 to P.K.C.).

## AUTHORS’ COMPETING INTERESTS STATEMENT

J.A.M. is cofounder and has an equity position in Arch Biopartners Inc. All other authors declare no conflicts of interest.

## CONTRIBUTORS’ STATEMENT

M.M. and N.A. designed, performed and analyzed the experiments. M.M. and J.A.M. wrote the manuscript; M.M., N.A. and J.A.M. prepared figures. P.C. extended access to the CLAMS metabolic cages and provided intellectual contributions to the project. J.A.M. supervised trainees and coordinated the study. All authors reviewed the results and approved the final version of the manuscript.

